# N-terminal mutations R5L and R5H share a common mechanism in disrupting tau condensates on microtubule surfaces

**DOI:** 10.64898/2026.11.13.766512

**Authors:** Rashmi Gangwar, Regina Agulto, Zack Quezada, Dai Zhai, Fabien Pinaud, Peter Chung

## Abstract

Tau, an intrinsically disordered protein abundant in neuronal axons, forms cooperative condensates on microtubule (MT) surfaces that regulate motor proteins and protect MTs from severing enzymes. Disease-associated mutations at residue 5 in the N-terminal projection domain of tau, far from the MT-tau binding interface, have been linked to progressive supranuclear palsy (PSP) (R5L) and frontotemporal dementia (FTD) (R5H). While R5L was previously shown to disrupt tau condensate formation on MT surfaces, it remained unclear whether R5H shares this phenotype. Using total internal reflection fluorescence (TIRF) microscopy, we show that both mutations reduce condensate packing density compared to WT tau under sub-saturating conditions at physiological pH (7.4), indicating impaired tau–tau cooperativity despite their distal location from the MT–tau binding interface. Single-molecule measurements further show both the dwell time and landing rate of tau within condensates are reduced while behavior on bare MTs remains unchanged, implicating weakened tau-tau cooperativity rather than disrupted binding to MTs. Strikingly, mutational analysis identifies the positive charge at residue 5 as critical for tau condensate formation: the charge-preserving R5K substitution retains WT-like behavior, whereas the charge-eliminating R5I substitution behaves like R5H/R5L. Further supporting this mechanism, R5H exhibits a pronounced pH dependence, transitioning from WT-like behavior at pH 6.4 to R5L-like behavior at pH 7.4 upon histidine deprotonation. Together, these findings establish that the positive charge at R5 is critical for tau condensate formation on MT surfaces and identify a common mechanism linking PSP and FTD.

**SIGNIFICANCE:** Tau, a protein that stabilizes microtubules in neurons, is central to Alzheimer’s disease and related neurodegenerative disorders. Many disease-associated tau mutations lie in its microtubule-binding region, but how mutations outside this region cause dysfunction remains unclear. We show that two such mutations, R5L (linked to progressive supranuclear palsy) and R5H (linked to frontotemporal dementia), weaken tau–tau interactions, so fewer tau molecules assemble together on microtubules, even though each molecule still binds microtubules with similar affinity. Restoring a positive charge at residue 5, either with another positively charged residue or by protonating R5H at lower pH, rescues assembly. These findings unite two mutations with distinct clinical presentations under a common molecular phenotype on microtubule surfaces.

## I. INTRODUCTION

Microtubules (MTs), one of the primary components of eukaryotic cytoskeletons, are protein nanotubes that are involved in a variety of cellular functions, including intracellular trafficking, cell motility, and chromosomal segregation[1–4]. This diversity of functions is mediated, in part, by microtubule-associated proteins (MAPs), which help regulate MT dynamics and structure. One such MAP, tau, is an intrinsically disordered protein primarily found in neuronal axons and, upon binding to MTs, suppresses their dynamic instability [5, 6]. Although tau is developmentally regulated in neurons and is critical for maintaining axonal health, tau dysfunction has been associated with many neurodegenerative diseases (tauopathies), including Alzheimer’s disease, frontotemporal dementia (FTD), and progressive supranuclear palsy (PSP)[7–12].

Due to alternative splicing of exons 2, 3, and 10, there are six isoforms of tau found in the human brain. Tau generally comprises a projection domain (with variable length, set by exons 2 and 3), a proline-rich region, a MT-binding region (containing either three or four imperfect MT-binding repeats, set by exon 10), and a C-terminal tail. Attention has often focused on the MT-binding region (and its flanking proline-rich region and C-terminal tail) for two reasons. First, these regions are known to directly bind to MTs and even facilitate distinct binding modes at physiological tau concentrations, including the recent discovery of larger-order complexes on MT surfaces (referred to as patches, cohesive islands, or condensates) that regulate MT interactions with motor proteins and MT-severing enzymes[4, 13–15]. Second, these regions contain the overwhelming bulk of the mutations and post-translational modifications of tau linked to disease. As a result, studies have often focused on how perturbations to these regions alter the interaction of tau with MTs[15–27].

Comparatively less is known regarding the role of molecular perturbations to the acidic projection domain of tau. While the length of the projection domain unequivocally controls inter-MT interactions[28, 29], mutations in the projection domain are relatively rare and their functional consequences remain poorly understood. Among the few classified as likely pathogenic is R5L, a missense mutation associated with PSP[30]. While R5L was shown to reduce MT assembly (through an unknown mechanism), recent work has shown R5L alters interactions between tau within tau condensates on MT surfaces[31]. Although full-length R5L tau (hereinafter referred to as R5L tau) exhibited similar binding affinity to MTs as full-length wild-type tau (hereinafter referred to as WT tau), the mutation disrupted the formation of tau condensates on MT surfaces at physiological tau concentrations. This is especially intriguing given that a different mutation at the exact same residue, R5H, is linked to FTD which exhibits some PSP-like pathology[32].

Here, we use a cell-free reconstitution with total internal reflection fluorescence (TIRF) microscopy to show that R5L and R5H tau similarly impair condensate formation on MT surfaces, reducing tau incorporation into condensates and tau density within condensates. Single-molecule tracking further reveals that both mutations decrease the landing rate and dwell time of tau within condensates, while leaving binding to bare MTs unchanged. Using mutational analysis, we show that the positive charge at residue 5 is critical for WT-like behavior. This mechanism is further corroborated by pH titration experiments, in which R5H tau behaves like WT at pH 6.4 (when His5 is protonated) but converges toward R5L at physiological pH 7.4. Together, these findings demonstrate that loss of positive charge at a single residue in the projection domain, a region of tau that does not directly bind MTs, is sufficient to disrupt the formation of higher-order tau structures on MT surfaces.

## II. RESULTS

### A. R5L and R5H reduce tau condensate formation on microtubules

To assess how the N-terminal disease-associated mutations R5H and R5L influence tau– microtubule interactions, we measured the binding of C-terminally sfGFP-tagged tau (WT, R5L, and R5H) to microtubules using TIRF microscopy. Taxol-stabilized, HiLyte 647-labeled MTs (100 nM) were immobilized on glass coverslips, with each tau variant introduced at increasing concentration, from 0.2 to 400 nM (Fig. S1). These experiments were performed in HEPES-based buffer at physiological pH (7.4) and salt concentration, as previously described[4, 14]. To establish consistency with prior observations for R5L in BRB80 buffer[31], we confirmed that the quantitative trends were similar across both buffers (Fig. S1). As previously reported, both WT and R5L tau variants formed discrete condensates on MT surfaces at concentrations as low as 2 nM (Fig. 1A and B)[14, 31]. We found that R5H tau also formed condensates under these conditions, establishing that the N-terminal mutation does not abolish higher-order assembly. Condensates of all the variants expanded and coalesced along the MT surface as the tau concentration increased from sub-saturating to saturating conditions (*≥* 200 nM) (Fig. 1A–C). However, condensate formation for both R5L and R5H tau appeared visibly reduced, decorating MTs more sparsely and discontinuously than WT.

**FIG. 1:**
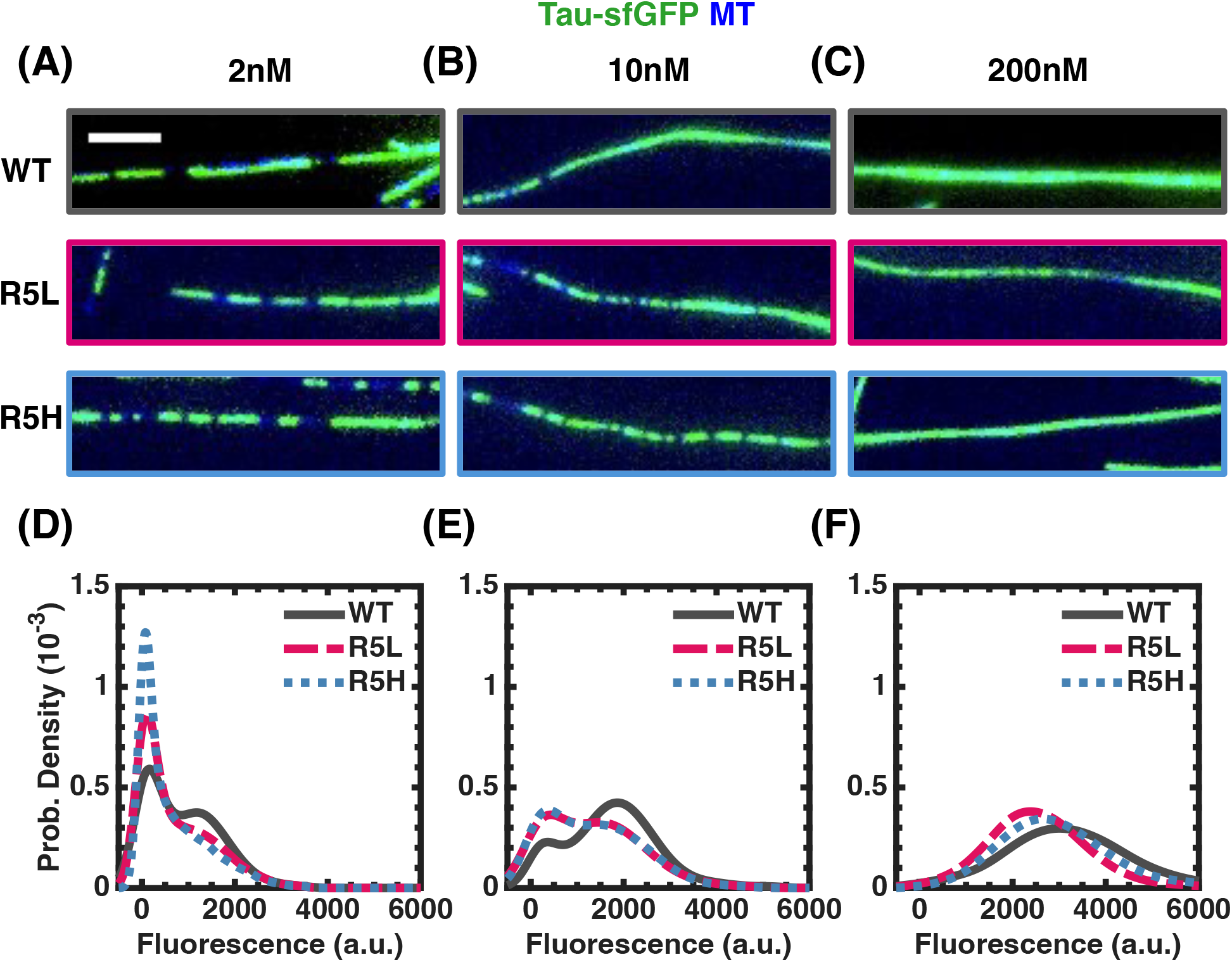
R5L and R5H reduce tau condensate formation on microtubules. (A–C) Representative fluorescence microscopy images showing condensate assembly along individual microtubules (blue) for WT, R5H, and R5L constructs (green) across increasing protein concentrations: **(A)** 2 nM, **(B)** 10 nM, and **(C)** 200 nM. Colored panel borders highlight construct-specific channels: WT (gray), R5L (ruby red), and R5H (cyan). Scale bar = 5 *µ*m. **(D–F)** Corresponding probability density distributions of tau fluorescence intensity (10*^−^*^3^) along the microtubule surface for **(D)** 2 nM, **(E)** 10 nM, and **(F)** 200 nM. Curves represent probability density estimates for WT (solid gray line), R5L (dash-dot ruby red line), and R5H (dotted cyan line). Data were pooled from *N* = 3 independent experiments comprising 15 acquisition areas, with each area containing *≈* 20–30 microtubules.

To evaluate this reduction in condensate formation, we plotted the probability density distributions of tau fluorescence intensity on MT surfaces (Fig. 1D-F and S2). At sub-saturating tau concentrations (Fig. 1D,E and S2), all variants exhibited bimodal intensity distributions that reflected the coexistence of diffusely bound tau (low-intensity peak) and the condensate state (high-intensity peak). However, both R5L and R5H tau exhibited stronger low-intensity peaks (and correspondingly weaker high-intensity peaks) compared to WT, suggesting both mutations reduce the propensity of tau to enter the condensate state. At saturating tau concentrations (200 nM, Fig. 1F), all variants converged into a unimodal, high-intensity peak, consistent with the diffuse-to-condensate transition and nearly complete coverage of the MT surface. Although mutations shift the balance between the diffuse and condensate states, they are insufficient to abolish condensation.

To resolve the partitioning of tau between the diffuse and condensate states, we fit each bimodal intensity distribution at sub-saturating concentrations to a two-component Gaussian mixture, extracting the peak position (corresponding to the tau concentration in each state) and the relative area weight of each population (Fig. S3). Consistent with a model in which the mutations impair tau–tau interactions, both R5L and R5H exhibited lower condensate peak intensities than WT at both sub-saturating concentrations (2 nM: 885 *±* 33 and 706 *±* 36 a.u. *µ*m*^−^*^1^ for R5L and R5H, respectively, vs. 1098 *±* 13 a.u. *µ*m*^−^*^1^ for WT; 10 nM: 1519 *±* 12 and 1521 *±* 15 a.u. vs. 1853 *±* 4 a.u. *µ*m*^−^*^1^), with the divergence between the two mutants becoming apparent at saturating concentration. In contrast, the diffuse peak varied only slightly across variants at 2 nM (60 to 95 a.u. *µ*m*^−^*^1^) and was essentially identical at 10 nM (*∼*235 a.u. *µ*m*^−^*^1^) (Fig. S3). Despite this similarity, the diffuse area increased for both mutations (2 nM: 35.6 *±* 1.0% and 36.3 *±* 1.2% for R5L and R5H, respectively, vs. 29.6 *±* 0.7% for WT; 10 nM: 21.8 *±* 0.6% and 22.2 *±* 0.7% vs. 13.1 *±* 0.2%). These fits show that both mutations shift tau toward the diffuse state and reduce the density of the condensate state in sub-saturating conditions.

At saturation (200 nM), all variants nearly covered the entire MT surface and the intensity distributions became unimodal. Together, these observations indicate that the measurement is now dominated by the condensate phase alone and the diffuse population is no longer resolved as a distinct state. This allows a single Gaussian fit to be interpreted as reporting directly on condensate packing density. Only R5L showed a pronounced reduction in condensate peak intensity (2477*±*5 a.u. *µ*m*^−^*^1^) compared to WT (3096*±*8 a.u. *µ*m*^−^*^1^). Although R5H showed an intermediate value (2758 *±* 8 a.u. *µ*m*^−^*^1^), binding isotherms indicate that the maximum binding capacity (*B*_max_) for R5H is comparable to WT, while R5L showed a consistent decrease in two different buffer conditions (Fig. S1). While the convergence to a single, unimodal distribution at saturation allows assessment of *B*_max_, ensemble measurements cannot resolve the kinetics within the diffuse and condensate phases and thus necessitate single-molecule resolution.

### B. R5L and R5H selectively weaken tau–tau cooperativity within condensates

To probe the kinetics of individual tau proteins within the condensate and on bare MT surfaces, we performed single-molecule tracking of Alexa Fluor 647–labeled SNAP-tau introduced either onto MTs decorated with sfGFP-tagged tau condensates of the matching variant, or onto bare MTs. Kymographs along the MT axis were used to track SNAP-tagged tau, allowing individual binding and unbinding events to be resolved as discrete streaks (Fig. 2A). The duration of each streak was extracted and compiled into a cumulative dwell-time distribution, which was fitted to a single-exponential function to obtain the characteristic dwell time *τ* for each variant (Fig. 2B). For all three variants, dwell times were longer within condensates than on bare MTs, suggesting that cooperative interactions among neighboring tau molecules promote tau residence within condensates (Fig. 2C and TABLE S1). Despite this overall increase, both R5L and R5H tau exhibited shorter dwell times (*τ* = 4.53 *±* 0.04 s and 4.55 *±* 0.05 s, respectively) within condensate than WT tau (*τ* = 5.37 *±* 0.05 s, whose dwell time was consistent with previously reported values [17]) (Fig. 2C). Strikingly, on bare MT surfaces the dwell times for all three tau variants were comparable (WT: 3.42 *±* 0.02 s; R5L: 3.28 *±* 0.03 s; R5H: 3.54 *±* 0.05 s), indicating that the reduced dwell time within condensates is specific to the condensate environment without substantially altering its interactions with the bare MT lattice (Fig. 2C).

**FIG. 2:**
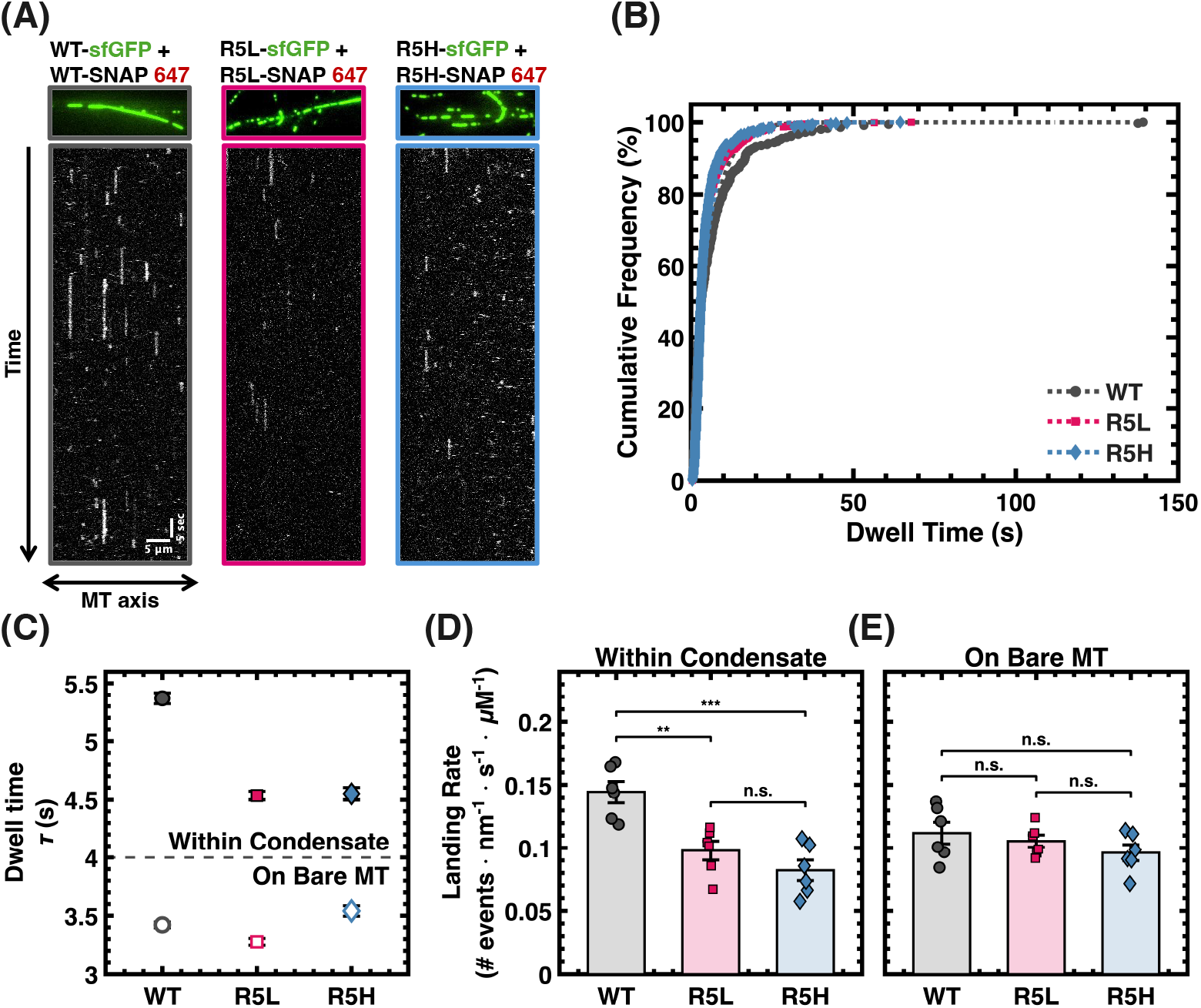
R5L and R5H selectively weaken tau–tau cooperativity within condensates. **(A)** Representative TIRF microscopy images (*top*) of sfGFP-tagged WT, R5L, and R5H tau (2 nM) forming condensates on MTs. Corresponding kymographs (*bottom*) show the trajectories of individual Alexa Fluor 647–labeled SNAP-tau molecules (50 pM) interacting within their respective condensates. Scale bars: 5 *µ*m (horizontal) and 5 s (vertical). **(B)** Cumulative dwell-time distributions of WT, R5L, and R5H tau molecules within their corresponding sfGFP-tau condensates. Dashed lines represent one-phase exponential fits used to determine the characteristic dwell time (*τ*). **(C)** Comparison of the characteristic dwell times (*τ*) of WT, R5L, and R5H tau within their respective condensates (solid markers) and on bare microtubules (hollow markers). Error bars represent standard error from fit. **(D-E)** Quantification of single-molecule landing rates of WT, R5L, and R5H tau within their respective **(D)** condensates, and **(E)** on bare MTs. Data are presented as mean *±* SEM from *N* = 6 independent experiments (*≈* 500 events). Statistical significance was evaluated using a pairwise post-hoc test (*∗p <* 0.05, *∗ ∗ p <* 0.01, *∗ ∗ ∗p <* 0.001; n.s., not significant).

Landing rate measurements confirm that changes to affinity are confined to the condensate; both R5L and R5H tau exhibited substantially reduced landing rates in condensates (0.098 *±* 0.008 and 0.082 *±* 0.008 nM*^−^*^1^s*^−^*^1^*µ*m*^−^*^1^ for R5L and R5H, respectively) compared to WT (0.144 *±* 0.008 nM*^−^*^1^s*^−^*^1^*µ*m*^−^*^1^) (Fig. 2D and TABLE S1), whereas the landing rates were statistically indistinguishable on bare MT surfaces (Fig. 2E and TABLE S1). This reduction in both dwell times and landing rates exclusively within the condensate is consistent with mutations diminishing tau–tau interactions as opposed to altering tau–MT surface interactions.

### C. Loss of charge at residue 5 impairs tau-tau cooperativity

To determine the molecular basis for the similarly impaired tau–tau cooperativity of the disease-associated R5L and R5H variants, despite their distinct side-chain properties (leucine is nonpolar, whereas histidine is classified as polar and basic but predominantly neutral at physiological pH), we produced two additional mutants: R5K, which retains the positive charge of arginine at residue 5, and R5I, which eliminates this charge with a nonpolar residue. As these new tau variants still formed condensates (Fig. 3A), we performed single-molecule measurements of SNAP-tagged tau within condensates of the matching tau variant and on bare MT surfaces to obtain dwell times and landing rates. Within condensates, R5K tau dwell times (*τ* = 5.77 *±* 0.06 *s*) and landing rates (0.163 *±* 0.009 nM*^−^*^1^s*^−^*^1^*µ*m*^−^*^1^) were comparable to WT tau, which had a slightly longer dwell time (Fig. 3B,C and TABLE S1). In contrast, R5I dwell times (*τ* = 4.01 *±* 0.05 *s*) and landing rates (0.095 *±* 0.007 nM*^−^*^1^s*^−^*^1^*µ*m*^−^*^1^) were reduced compared to WT tau and were comparable to the disease-associated mutants (TABLE S1). On bare MT surfaces, R5K tau dwell times (*τ* = 3.31 *±* 0.02 *s*) and landing rates (0.095 *±* 0.011 nM*^−^*^1^s*^−^*^1^*µ*m*^−^*^1^) were comparable to WT tau, whereas R5I exhibited a modest reduction in dwell time (*τ* = 3.01 *±* 0.02 *s*) but no significant change in landing rates (0.115 *±* 0.014 nM*^−^*^1^s*^−^*^1^*µ*m*^−^*^1^) (Fig. 3B,D and TABLE S1). While R5I exhibited a reduction in dwell time on bare MT surfaces, within condensates R5I showed a reduction of both dwell times and landing rates, indicating a stronger, condensate-specific effect. Taken together with the WT-like behavior of R5K and the R5L/R5H-like phenotype of R5I, these results indicate that loss of the positive charge at residue 5 impairs tau–tau cooperativity, potentially by weakening intermolecular electrostatic interactions that contribute to cooperative tau assembly on MT surfaces.

**FIG. 3:**
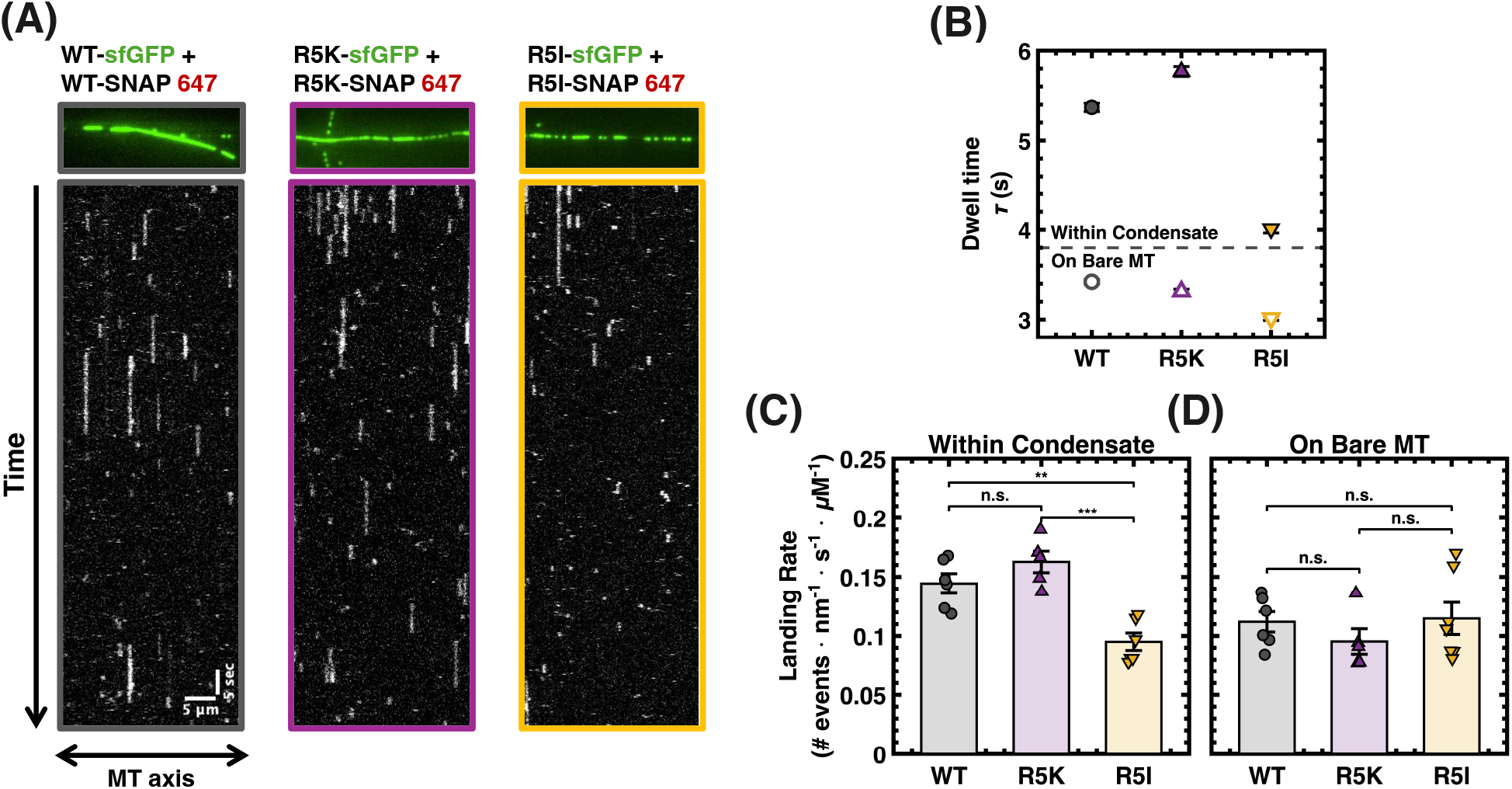
Loss of charge at residue 5 impairs tau-tau cooperativity. **(A)** Representative TIRF microscopy images (*top*) of sf-GFP tagged WT, R5K, and R5I tau (2 nM) forming condensates on MTs. Corresponding kymographs (*bottom*) show the trajectories of individual Alexa Fluor 647–labeled SNAP-tau molecules (50 pM) interacting within their respective condensates. Scale bars: 5 *µ*m (horizontal) and 5 s (vertical). **(B)** Comparison of the characteristic dwell times (*τ*) of WT, R5K, and R5I tau within their respective condensates (solid markers) and on bare microtubules (hollow markers). Error bars represent standard error from fit. **(C-D)** Quantification of single-molecule landing rates of WT, R5K, and R5I tau within their respective **(C)** condensates, and **(D)** on bare MTs. Data are presented as mean *±* SEM from *N* = 6 independent experiments (*≈* 500 events). Statistical significance was evaluated using a pairwise post-hoc test (*∗p <* 0.05, *∗ ∗ p <* 0.01, *∗ ∗ ∗p <* 0.001; n.s., not significant).

### D. Lower pH conditions partially rescue R5H tau condensate kinetics

While the similarity in condensate kinetics between R5L and R5H tau is striking, R5H possesses a unique biochemical property: unlike arginine, its protonation state can be tuned near physiological pH (the p*K*_a_ of histidine in model proteins has been found to be approximately 6.9, but can shift substantially based on sequence and local electrostatic environment[33]), allowing us to directly test whether reducing the pH can rescue tau–tau cooperativity within the condensate. We therefore performed single-molecule measurements of WT, R5L, and R5H tau across a range of physiological pH conditions (pH 6.4, 7.0, and 7.4) to test whether restoring the positive charge at residue 5 is sufficient to rescue the WT condensate phenotype. Dwell times for SNAP-tagged WT and R5L tau exhibited no substantial variation across the measured pH conditions within condensates of the same variant (Fig. 4A,B and TABLE I). However, dwell times for R5H tau decreased monotonically with increasing pH (*τ* = 5.91 *±* 0.07 s at pH 6.4 to *τ* = 4.55 *±* 0.05 s at pH 7.4) (Fig. 4C and TABLE I). Strikingly, at pH 7.4 the dwell times for both disease mutants were indistinguishable within condensates and at pH 6.4 the dwell times for R5H tau converged towards (and slightly exceeded) WT tau values (Fig. 4A–C and TABLE I). This pH-dependent rescue, specific to R5H and not R5L tau, is consistent with protonation of His5 at lower pH values restoring tau–tau cooperativity within the condensate.

**TABLE I:** Single-molecule dwell times, and landing rates of WT, R5L, and R5H tau within condensates across different pH conditions. Characteristic dwell times (*τ*) were obtained from one-phase exponential fits and are reported as *τ ±* standard error (SE) of the fit. Landing rates are reported as mean *±* SEM (*N* = 6).

|  | **WT** | **R5L** | **R5H** |
|---|---|---|---|
| **Dwell Time,** *τ* **(s)** |  |  |  |
| pH 6.4 | 5.51 ± 0.07 | 4.85 ± 0.06 | 5.91 ± 0.07 |
| pH 7.0 | 5.58 ± 0.06 | 4.82 ± 0.07 | 5.41 ± 0.06 |
| pH 7.4 | 5.37 ± 0.05 | 4.53 ± 0.04 | 4.55 ± 0.05 |
| **Landing Rate** (events · nm<sup>−1</sup> · s<sup>−1</sup> · *µ*M<sup>−1</sup>) |  |  |  |
| pH 6.4 | 0.209 ± 0.016 | 0.185 ± 0.013 | 0.241 ± 0.030 |
| pH 7.0 | 0.174 ± 0.009 | 0.155 ± 0.011 | 0.118 ± 0.009 |
| pH 7.4 | 0.144 ± 0.008 | 0.098 ± 0.008 | 0.082 ± 0.008 |

**FIG. 4:**
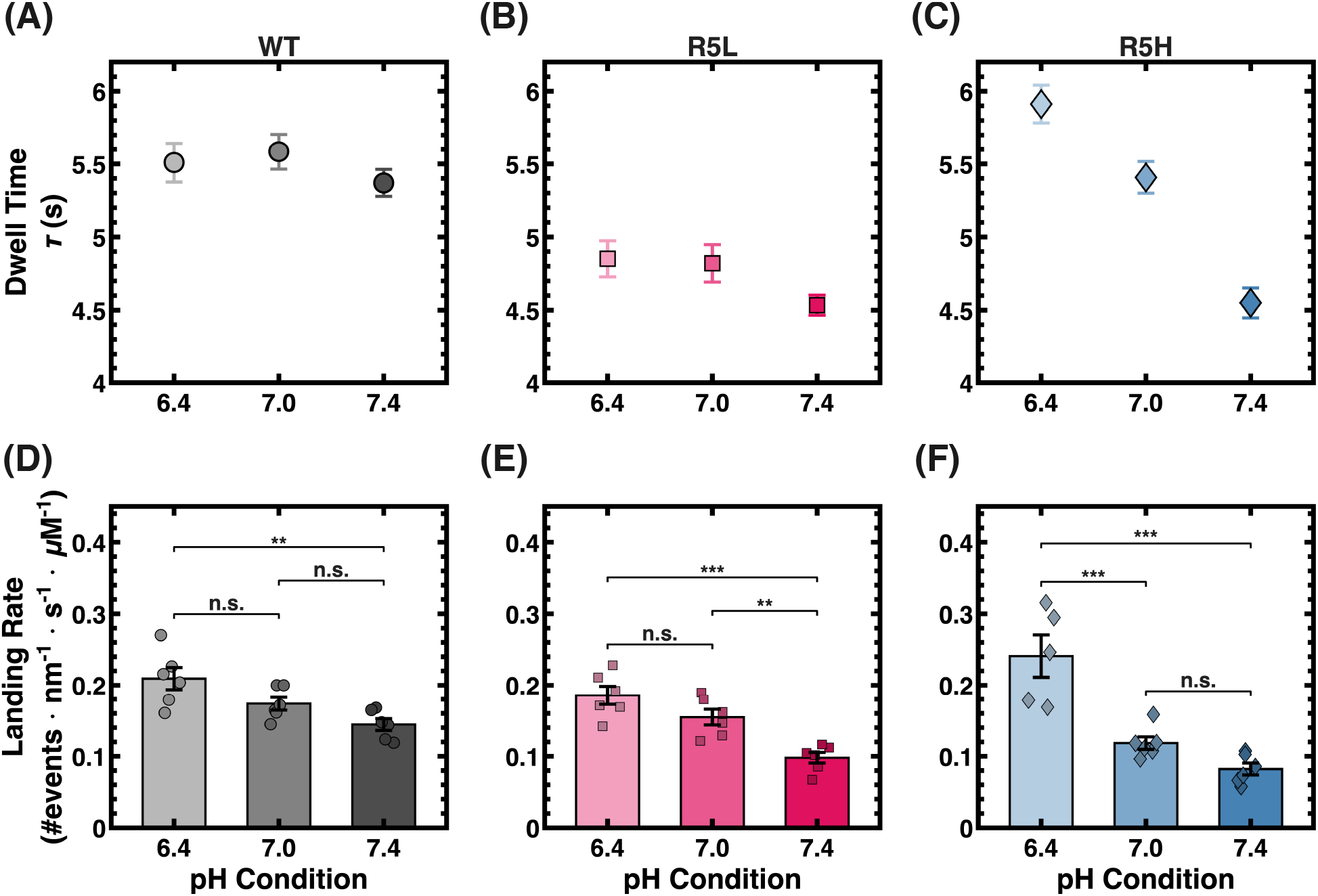
Lower pH conditions partially rescue R5H tau condensate kinetics. **(A–C)** Comparison of characteristic dwell times (*τ*) of single-molecule **(A)** WT, **(B)** R5L, and **(C)** R5H within their respective sfGFP-tau condensates on the microtubule surface across an increasing pH gradient. Error bars represent standard error from the fit. **(D–F)** Quantification of single-molecule landing rates for **(D)** WT, **(E)** R5L, and **(F)** R5H within their respective sfGFP-tau condensate regions. Data are presented as mean *±* SEM from *N* = 6 independent experiments (*≈* 500 events). Statistical significance was evaluated using a pairwise post-hoc test (*∗p <* 0.05, *∗ ∗ p <* 0.01, *∗ ∗ ∗p <* 0.001; n.s., not significant).

To ensure that this pH-dependent rescue in observed dwell times reflected a genuine recovery in the condensate phenotype for R5H, we examined the landing rates within condensates as a function of pH. We would expect a decrease in landing rates (due to the presence of histidines within the MT-binding region) with increasing pH, and all three variants exhibited this decrease (Fig. 4D–F and TABLE I). However, while the decrease in landing rates broadly tracked for WT and R5L tau, R5H tau exhibited a markedly steeper pH dependence: its landing rate within condensates exceeded that of WT tau at pH 6.4 (0.241 *±* 0.030 versus 0.209 *±* 0.016 nM*^−^*^1^s*^−^*^1^*µ*m*^−^*^1^) but fell well below both WT and R5L tau at pH 7.4 (0.082 *±* 0.008 versus 0.144 *±* 0.008 and 0.098 *±* 0.008 nM*^−^*^1^s*^−^*^1^*µ*m*^−^*^1^, respectively). Taken together, the convergence of R5H tau on WT tau-like behavior strongly supports a model in which positive charge at the fifth residue underlies tau–tau cooperativity within the condensate on MT surfaces.

## III. DISCUSSION

Although mutations and post-translational modifications that disrupt condensate formation have traditionally been mapped to the regions mediating multivalent intermolecular contacts, recent work has begun to reveal that domains distal to these regions can also modulate condensate behavior[34, 35]. Here, we demonstrate that the fifth residue of tau (far–removed from the MT-binding region and its flanking regions known to directly interact with MTs) is crucial not only for the formation of condensates on MT surfaces but also for tau–tau cooperativity within condensates. The completeness of the pH-dependent rescue of both single-molecule recruitment and retention underscores the sensitivity of condensate behavior to the charge state at this single position. This may be due, in part, to the geometry of tau bound to the MT surface: the projection domains of tau within condensates are likely densely packed, increasing the probability of contact between even distal residues and amplifying their contribution to intermolecular connectivity.

Although the R5L and R5H mutations occur at the same residue in tau, they have distinct clinical profiles: R5L is associated with progressive supranuclear palsy, whereas R5H is associated with FTD. Prior biochemical studies connecting the two mutations were limited to inter-protein interactions: both disrupt tau’s interaction with the dynactin complex and both alter tau–MT interactions[36, 37], though the mechanistic basis for the latter remains unclear. Here, we identify a shared phenotype based on interactions *between* tau, in a phenomenon that has been recently shown to directly control motor transport on MT surfaces and impair protection against severases[4, 13–15]. Indeed, their common mechanistic origin (as validated by our mutational analysis and pH experiments) has direct implications for another recently discovered mutation, R5C, that is associated with Parkinson’s disease[38]. Three mutations occurring at the same residue (each giving rise to distinct clinical presentations) raise the exciting possibility that the loss of charge at that residue leads to disruptions in condensate formation and impairment of tau–tau cooperativity, representing a common molecular mechanism that unites three distinct tauopathies.

Along these lines, future experiments should address the downstream impact of disruptions to tau condensate formation and decreased tau–tau cooperativity. Notably, residue 5 is located in the phosphatase-activating domain (PAD, residues 2–18) of tau, which has been shown to activate protein phosphatase 1*γ* (PP1*γ*) and initiate a signaling cascade that ultimately impairs fast axonal transport in many disease models[39–41]. Crucially, the R5L mutation has been shown to increase this impairment in an *ex vivo* axoplasm model, suggesting that condensate disruption at residue 5 may be upstream of aberrant PP1*γ* activation[42]. The resolution afforded by our cell-free system allows us to isolate the effect of a single mutation on collective tau behavior and generate hypotheses with predictive power in more complex biological systems.

## METHODS

### Microtubule assembly

Commercially available unlabeled (T240), biotinylated (T333P), HiLyte 647-labeled (TL670M), and HiLyte 488-labeled (TL488M) tubulin were purchased from Cytoskeleton, Inc, Denver, USA. For microtubule polymerization, 50 *µ*M unlabeled tubulin, 10 *µ*M biotinylated tubulin, and 5 *µ*M HiLyte 647- or HiLyte 488-labeled tubulin were mixed on ice in a total volume of 10 *µ*L of BRB80 buffer (80 mM PIPES, pH 6.8, 1 mM MgCl_2_, and 1 mM EGTA). GTP was added to a final concentration of 2 mM, and the mixture was incubated at 37*^◦^*C for 10–15 min to induce microtubule polymerization. The polymerized microtubules were further stabilized by the addition of 20 *µ*M taxol, followed by incubation at 37*^◦^*C for 20 min. To separate polymerized microtubules from unpolymerized tubulin, the samples were centrifuged through a 150 *µ*L 50% glycerol cushion prepared in BRB80 at 21,100 *×g* for 10 min at 37*^◦^*C. The resulting microtubule pellet was resuspended in 50 *µ*L BRB80T (BRB80 + 10 *µ*M taxol) buffer and used as the microtubule stock for subsequent experiments.

### Protein constructs

The pRBC-WT plasmid encoding the human full-length tau isoform 2N4R (441 amino acids) was purchased from Twist Bioscience, CA, USA. Fragments containing sfGFP, or SNAP-tag, followed by a linker GGS, TEV protease cleavage site (ENLYFQS) and a His_6_ tag, were cloned into the pRBC vector at the C terminus of rau using Gibson assembly, resulting in the following construct architecture: rau–GGS–sfGFP/SNAP–TEV–His_6_. Missense mutations at residue 5 of tau were introduced by site-directed mutagenesis using the NEB Site-Directed Mutagenesis Kit (New England Biolabs, Ipswich, MA) and primers synthesized by Genewiz, NJ, USA. The resulting plasmid constructs were sequence-verified by Plasmidsaurus, Inc, LA, CA.

### Expression and Purification of Tau

Tau plasmids were transformed into chemically competent *Escherichia coli* BL21(DE3) cells using a standard transformation protocol and plated on LB agar containing ampicillin. Following overnight incubation, several individual colonies were inoculated into 10 mL LB media containing ampicillin and grown overnight. The saturated overnight culture was subsequently used to inoculate 1 L of Terrific Broth (TB) media. Cells were grown at 37*^◦^*C with shaking at 250 rpm until the optical density at 600 nm (OD_600_) reached approximately 1.0. Protein expression was then induced with 0.5 mM IPTG, and the cultures were incubated at 21*^◦^*C with shaking at 250 rpm for 16–18 h.

Following protein expression, cells were harvested by centrifugation at 4,000 *×g* for 20 min at 4*^◦^*C. The cell pellet was resuspended in Ni-NTA lysis buffer (10 mM Tris, 5 mM imidazole, 500 mM NaCl, and 1 mM phenylmethylsulfonyl fluoride (PMSF), pH 8.0) and lysed by sonication for 10 min (2 s on, 2 s off, 50% amplitude) using a Qsonica Q125 sonicator (Qsonica, Newtown, CT, USA). The soluble fraction was separated from the cell debris by centrifugation at 72,000 *×g* for 30 min at 4*^◦^*C. The clarified lysate was incubated with Ni-NTA agarose resin for 60 min at 4*^◦^*C with continuous rotation. Following binding, the protein was eluted using Ni-NTA elution buffer (10 mM Tris, 300 mM imidazole, and 500 mM NaCl, pH 8.0).

The eluted protein was subsequently precipitated by the addition of ammonium sulfate [(NH_4_)_2_SO_4_] to 0.4 g/mL and centrifuged at 72,000 *×g* for 20 min at 4*^◦^*C. The resulting protein pellet was resuspended in super TEV cleavage buffer (50 mM Tris, 1 mM EDTA, 1 mM PMSF, and 1 mM Tris(2-carboxyethyl)phosphine (TCEP))[43]. TEV protease protein was added at 2.5% (w/w) relative to the mass of tau protein, and the sample was incubated overnight at 4*^◦^*C protected from light to cleave the C-terminal His_6_ tag. Following TEV cleavage, the sample was subjected to a second Ni-NTA affinity purification step, and the flow-through containing the His_6_-tag-cleaved tau protein was collected.

For final purification and buffer exchange, the protein was subjected to size-exclusion chromatography using a Superdex 200 Increase 10/300 GL column on an AKTA Pure system (Cytiva, Marlborough, MA, USA), equilibrated with BRB80 (sfGFP-tagged tau)/PBS saline (SNAP-Tagged tau) supplemented with 1 mM TCEP. Fractions corresponding to the tau protein peak were collected and pooled. The purified protein was aliquoted, flash-frozen, and stored at *−*80*^◦^*C until further use.

### Labeling tau proteins with SNAP tag substrate

Purified SNAP-tagged tau proteins were fluorescently labeled with SNAP-Surface^®^ Alexa Fluor^®^ 647 according to the manufacturer’s recommended protocol (New England Biolabs, Ipswich, MA). Briefly, 10–15 *µ*M purified SNAP-tagged tau was incubated with 20–30 *µ*M SNAP-Surface Alexa Fluor 647 substrate, corresponding to an approximately two-fold molar excess of dye, in phosphate-buffered saline (PBS; 137 mM NaCl, 2.7 mM KCl, 10 mM Na_2_HPO_4_, and 1.8 mM KH_2_PO_4_, pH 7.4). The labeling reaction was incubated at 37*^◦^*C for 30 min, followed by overnight incubation at 4*^◦^*C.

To remove unconjugated SNAP-Surface Alexa Fluor 647 and exchange the protein into the experimental buffer, the labeling reaction was run through a HiTrap Desalting 5 mL column (Cytiva) using an AKTA Pure system equilibrated with BRB80 supplemented with 1 mM TCEP. Fractions containing the labeled protein were collected, and the labeling efficiency was determined by UV–Vis spectroscopy using a NanoDrop spectrophotometer from the absorbance at 280 and 647 nm and the corresponding extinction coefficients. Labeling efficiencies of approximately 65–68% were routinely obtained. The labeled protein was aliquoted, flash-frozen, and stored at *−*80*^◦^*C until further use.

### TIRF microscopy assays

All TIRF microscopy experiments were performed at room temperature on an objective-based TIRF microscope equipped with a Nikon Eclipse Ti stand and a 100*×*/1.49 NA Apo TIRF objective (Nikon). Flow chambers were assembled using acid-washed glass coverslips, prepared as previously described [14, 17], pre-cleaned glass slides, and removable double-sided adhesive tape.

On the day of the experiment, flow chambers were incubated with 0.5 mg/mL PLL-PEG-biotin [PLL(20)-g[3.5]-PEG(2)/PEG(3.4)-biotin (50%); SuSoS Surface Technology, Switzerland] for 10–15 min, followed by washing with BRB80 buffer. The chambers were subsequently incubated with 0.5 mg/mL streptavidin for 5 min and washed again with BRB80. Taxol-stabilized microtubule stocks were diluted to approximately 100 nM in BRB80T. Diluted microtubules were flowed into the chamber and incubated for 5–10 min for adhesion on the chamber surface. Unbound microtubules were removed by washing the chambers with two chamber volumes of assay buffer (60 mM HEPES, 50 mM potassium acetate, 2 mM MgCl_2_, 1 mM EGTA, 0.5% Pluronic F-127, 0.1 mg/mL biotin-BSA, 0.2 mg/mL *κ*-casein, and 10 *µ*M taxol, pH 7.4). Tau proteins were diluted to the indicated concentrations in assay buffer and introduced into the flow chambers containing immobilized microtubules. The samples were incubated for at least 5 min before imaging. For single-molecule experiments, the assay buffer was supplemented with an oxygen-scavenging system consisting of Trolox/PCA/PCD (2 mM Trolox, 2 mM protocatechuic acid (PCA), and *∼*50 nM protocatechuate-3,4-dioxygenase (PCD)) to minimize fluorophore photobleaching during imaging.

### Single molecule experiments

For single-molecule measurements within tau high-density assemblies, microtubules were immobilized on the chamber surface as described above. SNAP-tagged tau labeled with Alexa Fluor 647 (50 pM) was mixed with sfGFP-tagged tau (2 nM) in assay buffer supplemented with the PCA/PCD/Trolox oxygen-scavenging system and introduced into the microtubule-containing flow chamber. Following incubation, time-lapse movies were acquired to monitor single-molecule tau dynamics along the microtubule lattice. Images were acquired at a frame interval of Δ*t* = 0.2 s for a total duration of 2 min 20 s.

For single-molecule measurements on the bare microtubule lattice, HiLyte 488-labeled microtubules were immobilized on the chamber surface. Alexa Fluor 647-labeled SNAP-tau was diluted to 20 pM in assay buffer supplemented with the PCA/PCD/Trolox oxygen-scavenging system and introduced into the chamber without the addition of unlabeled or sfGFP-tagged tau. Time-lapse images were acquired at a frame interval of Δ*t* = 0.1 s for a total duration of 1 min 10 s.

### Quantification of Tau fluorescence intensity along microtubules

Fluorescence microscopy images were visualized using Fiji/ImageJ (version 1.54s), and quantitative image analysis was performed using a custom MATLAB pipeline (MATLAB R2025b; MathWorks). Dual-channel fluorescence images consisted of a tau channel (*ch*_1_) and a microtubule channel (*ch*_2_). Microtubules were identified and segmented exclusively from the microtubule channel, and the resulting masks and skeleton coordinates were used to quantify tau fluorescence in the corresponding tau channel. Regions containing microtubule aggregates or other imaging artifacts were manually excluded when necessary.

#### Microtubule segmentation and tracing

Microtubule images were smoothed using a Gaussian filter with *σ* = 1.5 pixels. Microtubules were segmented by combining adaptive local thresholding (sensitivity = 0.5) with a global intensity cutoff defined as 1.2 times the median image intensity. Objects smaller than 50 pixels^2^ were removed, and holes within the resulting binary masks were filled. The refined binary masks were skeletonized to generate one-pixel-wide representations of individual microtubule structures. Branch points were identified and removed using a 3-pixel-radius morphological disk to minimize ambiguities arising from intersections and branching structures. The remaining skeletons were separated into individual linear segments, and only segments longer than 0.5 µm were retained for analysis. Images were calibrated at 6.25 pixels per micrometer, corresponding to a spatial sampling interval of approximately 0.16 µm per pixel.

#### Tau fluorescence intensity and local background correction

For each retained microtubule segment, the skeleton was dilated using a 2-pixel-radius morphological disk to generate a region representing the microtubule width. The mean tau fluorescence within this region was calculated from the tau channel. Local background fluorescence was estimated from a surrounding region generated by dilating the microtubule mask by 15 pixels and excluding both the microtubule mask and neighboring microtubule structures. The median fluorescence intensity within this local background region was used as the background intensity. The net tau fluorescence intensity was calculated as

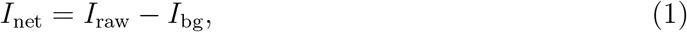

where *I*_raw_ is the mean tau fluorescence within the microtubule segment mask and *I*_bg_ is the median fluorescence within the surrounding local background region. Segments with insufficient raw signal or net intensity were excluded from subsequent analysis.

To obtain spatially resolved tau intensity profiles along individual microtubules, the endpoints of each skeleton segment were identified and the skeleton was traced sequentially to generate ordered coordinates. The cumulative distance along each traced segment was calculated and converted from pixels to micrometers. At each skeleton position, tau fluorescence was measured as the mean intensity within a 3 *×* 3-pixel neighborhood centered on the corresponding skeleton coordinate. The same local background value determined for the corresponding microtubule segment was subtracted from each pointwise measurement to obtain the net tau intensity profile as a function of distance along the microtubule.

#### Tau intensity distributions

For each construct, net tau fluorescence measurements collected from all analyzed microtubule segments were pooled to generate intensity distributions. Histograms were calculated using 50 intensity bins, and bin counts were normalized by the total analyzed microtubule length to report the frequency of tau intensity measurements per unit microtubule length.

Continuous probability density functions were generated using Gaussian kernel density estimation with MATLAB’s ksdensity function. For comparison between constructs, intensity distributions were evaluated on a common grid of 200 equally spaced intensity values spanning the combined intensity range of the datasets. The probability density was calculated as

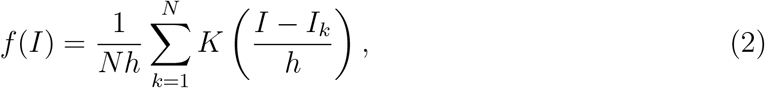

where *N* is the total number of intensity measurements, *I_k_* is the net tau fluorescence intensity of the *k*th measurement, *I* is the intensity at which the probability density is evaluated, *K* is the Gaussian kernel function, and *h* is the bandwidth determined by ksdensity.

### Single molecule analysis

Single-molecule tau dynamics were analyzed using kymographs generated in Fiji/ImageJ with the Kymograph plugin. Kymographs were generated by manually tracing a line along the microtubule lattice. For measurements within high-density tau assemblies, the sfGFP-tau channel was used as a fiducial marker to identify the boundaries of high-density and bare microtubule regions. Individual Alexa Fluor 647-labeled SNAP-tau binding events were identified from the corresponding single-molecule channel, and their residence times were determined from the duration of each event in the kymograph. Only events for which the complete binding trajectory, from initial association to dissociation from the microtubule, was captured within the acquisition window were included in the dwell-time analysis. Events longer than three pixels were selected as binding events for analysis. The same procedure was used for bare-microtubule measurements, with kymographs generated directly along the microtubule lattice.

For each condition, the cumulative frequency of binding events was plotted as a function of dwell time and fitted to a single-exponential association function using a custom MATLAB routine:

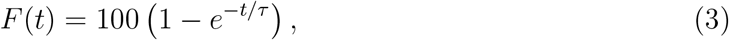

where *F* (*t*) is the relative cumulative frequency of binding events at time *t*, and *τ* is the fitted characteristic dwell time.

Tau landing rates were determined by counting individual Alexa Fluor 647-labeled SNAP-tau binding events within the condensate on microtubule lattice. The number of landing events was normalized by the total analyzed microtubule length, acquisition time, and concentration of the fluorescently labeled tau:

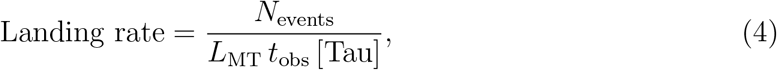

where *N*_events_ is the number of observed tau-binding events, *L*_MT_ is the analyzed microtubule length, *t*_obs_ is the observation time, and [Tau] is the concentration of Alexa Fluor 647-labeled SNAP-tau.

### Statistics and reproducibility

Details of statistical analyses, sample sizes, and the number of independent experiments are provided in the corresponding figure captions.

## Code availability

Data analysis was performed using custom-written MATLAB scripts. The MATLAB code used to analyze the data presented in this study is available from the corresponding authors upon request.

