## Supplementary Information file for "N-terminal mutations R5L and R5H share a common mechanism in disrupting tau condensates on microtubule surfaces"

Peter Chung

*Department of Physics & Astronomy, Department of Chemistry,*

*Mann Department of Biomedical Engineering,*

*University of Southern California, Los Angeles, California 90089, USA*

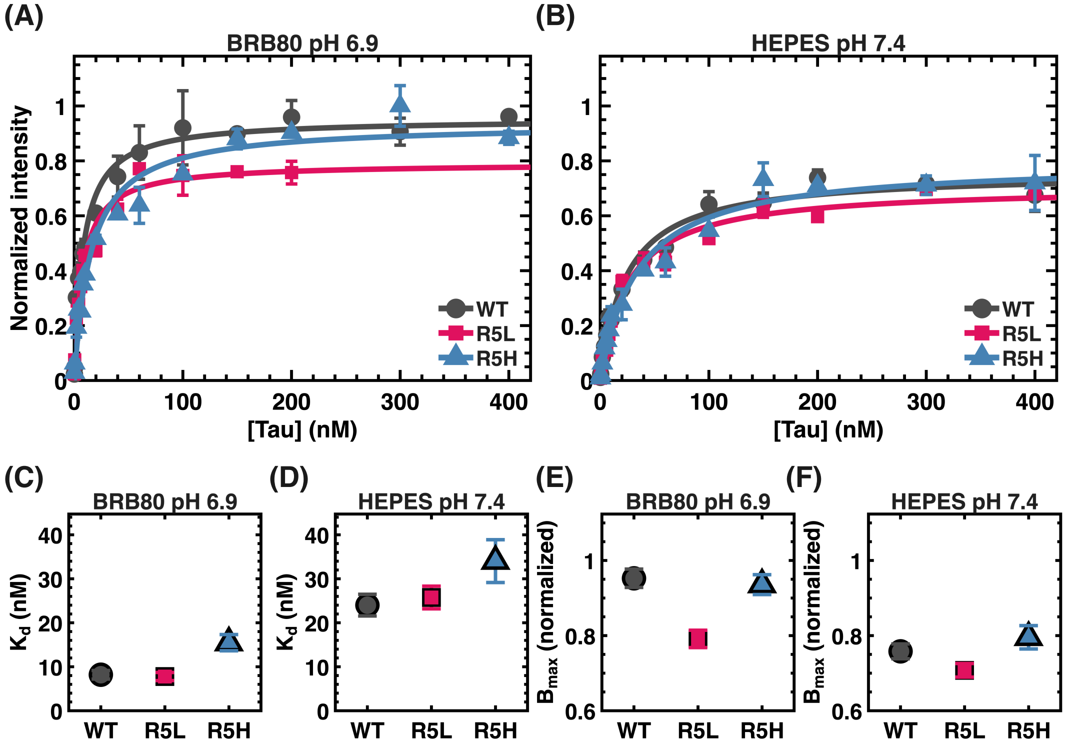


FIG. S1. **Equilibrium binding curves of WT, R5L, and R5H tau to MTs. (A, B)** Binding isotherms of sfGFP-tau WT (gray circles), R5L (magenta squares), and R5H (blue triangles) on MTs in (A) BRB80 (80 mM PIPES, 1 mM $MgCl_2$, 1 mM EGTA, pH 6.9) and (B) HEPES-based buffer (60 mM HEPES, 50 mM potassium acetate, 2 mM $MgCl_2$, 1 mM EGTA, pH 7.4). Normalized tau fluorescence intensity on MTs is plotted as a function of tau concentration (0.2–400 nM). Solid lines show fits to a Hill's equation. **(C, D)** Dissociation constants ($K_d$) obtained from the fits in (A) and (B), respectively. **(E, F)** Maximal binding capacity ($B_{\text{max}}$) obtained from the fits in (A) and (B), respectively. Data are presented as mean ± SEM; error bars in (C–F) represent the standard error of the fit.

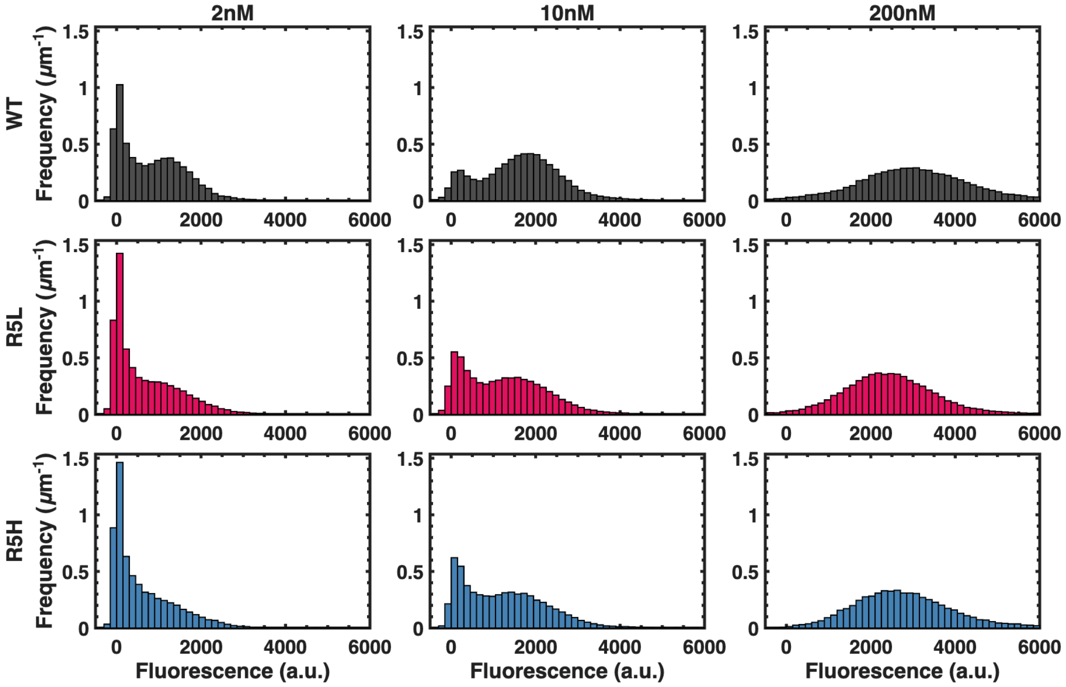


FIG. S2. **Fluorescence intensity distributions of WT, R5L, and R5H tau on MT surfaces.** Histograms of sfGFP-tau fluorescence intensity along MTs for WT (gray, top row), R5L (ruby red, middle row), and R5H (cyan, bottom row) tau at 2 nM (left), 10 nM (middle), and 200 nM (right). At subsaturating concentrations (2 and 10 nM), the distributions are bimodal, with a low-intensity diffuse population and a high-intensity condensate population. At saturation (200 nM), the distributions become unimodal, reflecting near-complete coverage of the MT surface by condensates. Data were pooled from $N = 3$ independent experiments comprising 15 acquisition areas, with each area containing ≈ 20–30 microtubules; bin width, 50 a.u.

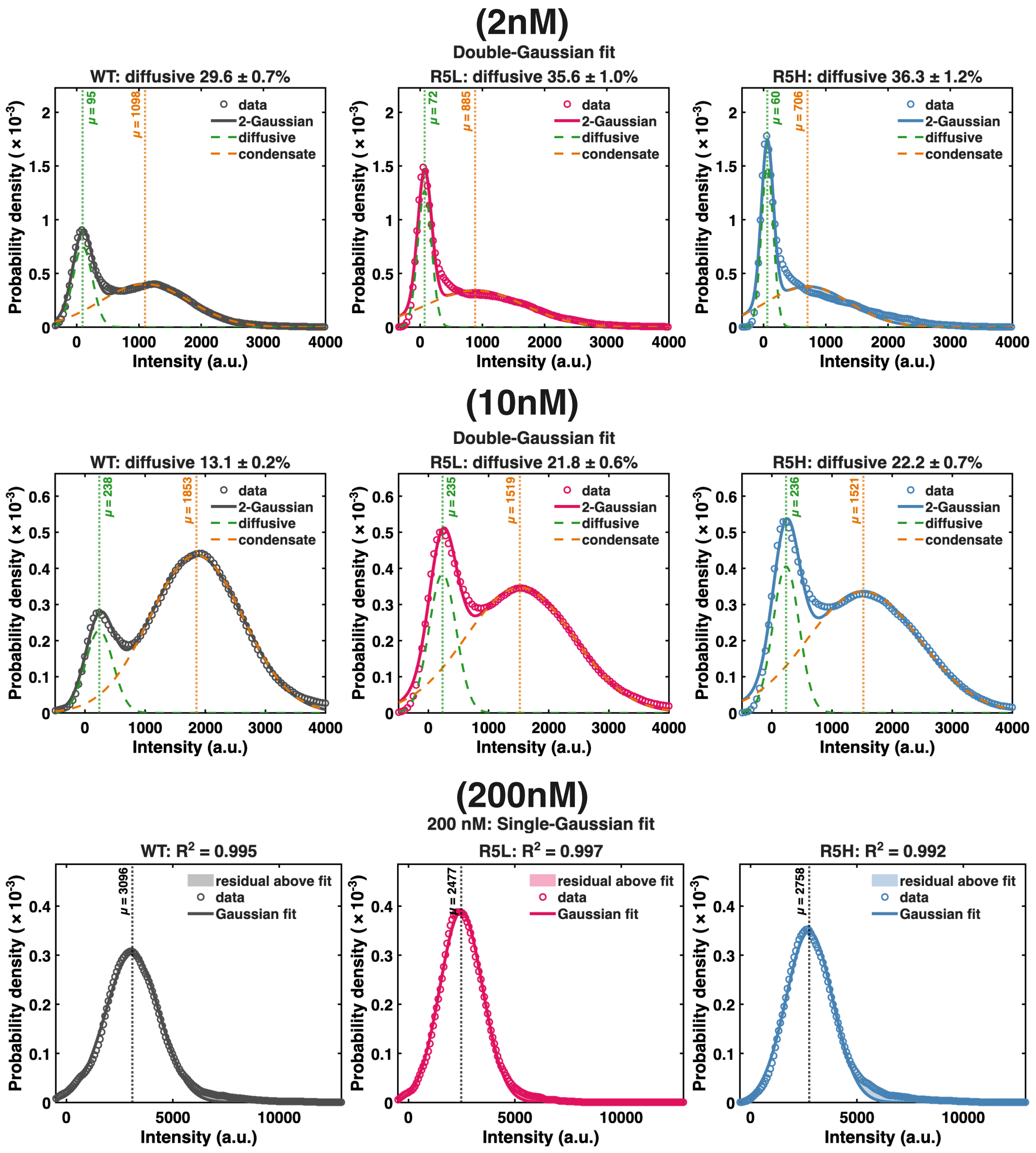


FIG. S3. **Gaussian mixture analysis of tau intensity distributions on MT surfaces.** Probability density of sfGFP-tau fluorescence intensity on MTs for WT (gray, left), R5L (ruby red, middle), and R5H (cyan, right) tau. **(Top and middle rows)** At subsaturating concentrations (2 and 10 nM), distributions were fit to a two-component Gaussian mixture (solid line), resolving a low-intensity diffuse population (green dashed line) and a high-intensity condensate population (orange dashed line). Vertical dotted lines mark the fitted peak positions ($\mu$), and panel titles give the area fraction of the diffuse population. **(Bottom row)** At saturation (200 nM), distributions were unimodal and were fit to a single Gaussian; the fitted peak position ($\mu$) and goodness of fit ($R^2$) are indicated. Shaded regions show the residual of the data above the fit. Values are reported as mean ± SE.

TABLE S1. Characteristic dwell times and single-molecule landing rates of WT and tau variants. Characteristic dwell times ($\tau$) were obtained from one-phase exponential fits and are reported as $\tau \pm$ standard error (SE) of the fit. Landing rates within condensates and on bare microtubules (MTs) are presented as mean $\pm$ SEM. Dwell time and landing rates are pooled from $N = 6$ experiments, $\approx n = 500$ events

| | **Dwell Time** $\tau$ (s) | | **Landing Rate** (events $\cdot$ nm$^{-1} \cdot$ s$^{-1} \cdot \mu$M$^{-1}$) | |
|---|---|---|---|---|
| **Construct** | **Within Condensates** | **On Bare MTs** | **Within Condensates** | **On Bare MTs** |
| WT | $5.37 \pm 0.05$ | $3.42 \pm 0.02$ | $0.144 \pm 0.008$ | $0.112 \pm 0.009$ |
| R5H | $4.55 \pm 0.05$ | $3.54 \pm 0.05$ | $0.082 \pm 0.008$ | $0.096 \pm 0.006$ |
| R5L | $4.53 \pm 0.04$ | $3.28 \pm 0.03$ | $0.098 \pm 0.008$ | $0.105 \pm 0.005$ |
| R5K | $5.77 \pm 0.06$ | $3.31 \pm 0.02$ | $0.163 \pm 0.009$ | $0.095 \pm 0.011$ |
| R5I | $4.01 \pm 0.05$ | $3.01 \pm 0.02$ | $0.095 \pm 0.007$ | $0.115 \pm 0.014$ |

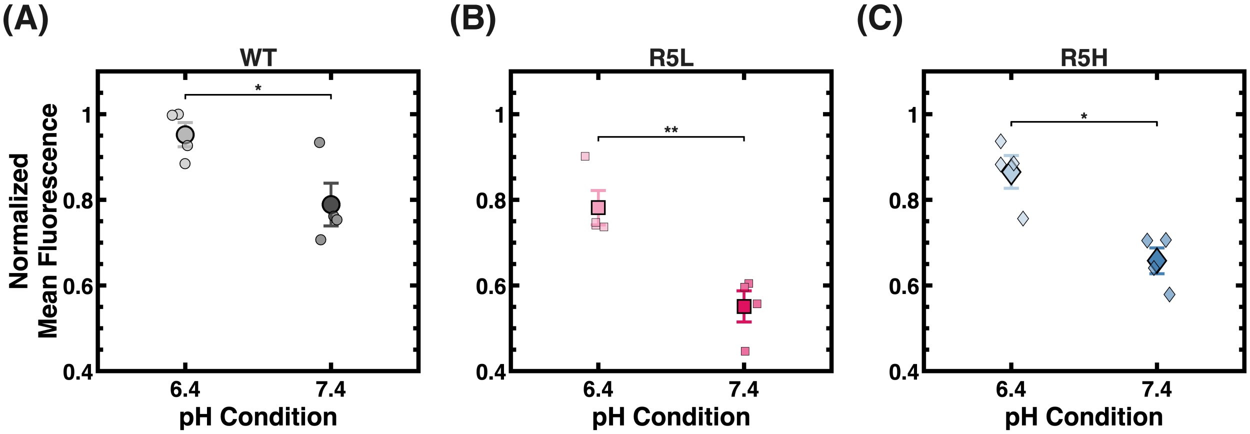


FIG. S4. **(A–C)** Normalized mean fluorescence intensity of 5 nM sfGFP-tau variants **(A)** WT, **(B)** R5L, and **(C)** R5H on the microtubule surface at (HEPES-based buffer) pH 6.4 and 7.4, mean $\pm$ SEM, from $N = 4$ independent experiments. Statistical significance was evaluated using a pairwise post-hoc test ($*p < 0.05$, $**p < 0.01$, $***p < 0.001$; n.s., not significant).
